# Evaluation of Cognitive Function in the Dog Aging Project: Associations with Baseline Canine Characteristics

**DOI:** 10.1101/2022.05.04.490636

**Authors:** Sarah Yarborough, Annette Fitzpatrick, Stephen M. Schwartz, Dog Aging Project Consortium

## Abstract

Canine Cognitive Dysfunction (CCD) is a neurodegenerative disease in aging dogs. It has been described previously in relatively small cohorts of dogs using multiple different rating scales. This study aimed to use a minimally modified CCD rating scale developed by previous researchers to describe the prevalence of CCD more thoroughly in a large, nationwide cohort of companion dogs participating in the Dog Aging Project (DAP). Associations between various canine characteristics, predicted lifespan quartiles, and CCD were examined using univariable and multivariable logistic regression models and Receiver Operating Curve (ROC) analysis.

When controlling for all other characteristics, the odds of CCD increased 52% with each additional year of age. Among dogs of the same age, health status, breed type, and sterilization status, odds of CCD were 6.47 times higher in dogs who were not active compared to those who were very active. When controlling for age, breed type, activity level, and other comorbidities, dogs with a history of neurological, eye, or ear disorders had higher odds of CCD. Lifespan quartile analysis showed excellent discriminating ability between CCD positive and negative dogs. Weight-based lifespan quartile estimation could therefore serve as a tool to inform CCD screening by veterinarians.

## Introduction

Neurodegenerative diseases associated with aging have become increasingly prevalent among the aging American population. In 2020, approximately 5.8 million Americans were living with Alzheimer’s disease (AD). It is the sixth leading cause of death in the country^1^. Despite this, the complex pathological pathways that lead to the development of AD are still not fully understood and can be difficult to study *in vivo* in early phases of disease progression. While transgenic animal models have been extensively used to study the pathophysiology of AD, limitations have been identified that have prompted investigation into non-transgenic animal models^2^.

The clinical and histological presentation of human AD and Canine Cognitive Dysfunction (CCD) in dogs have many similarities. As with humans, canine cognitive function declines throughout the course of the dog’s lifespan^3^. Clinical signs of this decline appear to be related to learning and memory deficits, loss of spatial awareness, altered social interactions, and disrupted sleeping patterns^4,5,6^. Further, the presentation of human AD and CCD share certain neuropathological features such as amyloid-β plaque deposition^2,7,8,9,10^.

The observed parallels between CCD and human AD suggest that dogs exhibiting CCD may offer researchers a valuable animal model in which to study characteristics of neurodegenerative diseases that are relevant to, but challenging to study in, human populations^11^. Further, dogs with CCD could serve as candidates for AD preventative and/or therapeutic strategies^2^. Finally, CCD is also a major health concern of dog owners and veterinarians. An increased understanding of CCD may help to advance treatment of cognitive disease in dogs. These potential benefits highlight the importance of accurate diagnosis of CCD in companion dogs.

A wide range of structured interviews and theory-driven scales for evaluating CCD have been developed within the last 20 years^4,12,13,14^. Studies of these scales have shown a correspondingly wide range of prevalence estimates that nonetheless appear to indicate an increased prevalence of cognitive impairment as a dog ages^7,14,15^. In order to gain a better understanding of CCD in aging dogs, researchers recently have developed assessment tools that are able to distinguish cognitively impaired aged dogs from those who are experiencing healthy aging^11,14^.

These scales provide veterinarians and researchers with tools to better identify the prevalence of CCD in aging dogs but have only been used to describe cognitive function in relatively small populations (<1,000 dogs)^11,14^. The purpose of this study was to describe the range of cognitive function scores and prevalence of CCD in a very large, nationwide sample of companion dogs participating in the Dog Aging Project (DAP) (n = 27,542). The DAP utilized the Canine Social and Learned Behavior (CSLB) survey, a minimally modified version of the validated canine cognitive dysfunction rating scale (CCDR) developed by Salvin *et al*.^4^. Further, we examined associations between weight-based lifespan quartile and CCD. Classifying a dog’s lifespan into quartiles and quantifying the predictive ability of the CCDR in each quartile potentially allows for preventative healthcare measures to be taken by owners and veterinarians. The results could eventually lead to more timely detection and treatment of CCD^16^.

## Methods

### Study Setting

Data were obtained from the DAP, a nationwide longitudinal study on aging initiated in 2018. One goal of the DAP is to better define aging in dogs based on comorbidities, frailty, and inflammaging^17^. Companion dog owners interested in enrolling their dogs complete multiple surveys throughout their involvement in the DAP. For the purposes of this report, these included the baseline Health and Life Experience Survey (HLES) and the CSLB survey completed during the first-year enrollment in DAP. The HLES is an extensive online survey that is distributed to all dog owners upon baseline enrollment, with sections on dog and owner household demographic characteristics, dog physical activity, environment, behavior, diet, medications and preventatives, and health status. The CSLB survey is a 13-item questionnaire based on Salvin *et al*.’s CCDR scale that aims to assess CCD^4^. It is described in further detail below.

### Study Design

This study analyzed baseline associations between selected characteristics collected through the HLES and dog cognitive characteristics reported in the CSLB. HLES data were provided by participants immediately upon enrollment, and CSLB data were provided by participants when that survey was administered. Dates of collection for HLES data used in the study ranged from December 23, 2019, to December 11, 2020. Dates of collection for CSLB data used in the study ranged from October 30, 2020, to December 31, 2020. The median time between completion of the two surveys was 5 weeks (range: 0–50.3 weeks).

### Study Subjects

All dogs whose owners did not complete both the baseline HLES and CSLB surveys prior to December 31, 2020, were excluded from study. Further, all dogs whose owners indicated that they were not able to confidently report their dog’s age, as assessed in the HLES, were also excluded. We began with a study sample of 27,542 dogs whose owners had completed the baseline HLES; 20,096 of those dogs’ owners had also completed the CSLB survey. Of those 20,096 dogs, 15,019 owners reported being certain of their dog’s age. The final sample for this study consisted of 15,019 dogs.

### Data Sources

The validated CCDR instrument, used by Salvin *et al*.^*4*^ for the identification of CCD, was minimally modified and renamed the CSLB (Canine Social and Learned Behavior Survey) for this study. In contrast to the original CCDR: 1) the instrument name was modified so as not to suggest dementia or cognitive dysfunction (identified in the term CCD) to the owners of enrolled dogs; and 2) it was programmed as an on-line digital tool with American English spelling. The CSLB thus consisted of 13 questions that assessed behaviors such as getting stuck behind objects, pacing, and failing to recognize familiar people. Responses for each question were scored based on frequency of the behavior (1=never, 2=once a month, 3=once a week, 4=once a day, 5=more than once a day). In addition, some questions asked owners to compare the severity of their dog’s current behavior to the severity of the behavior six months prior. Numeric scores were summed from each owner’s responses. Certain questions had multipliers based on their clinical severity, as identified in CCD diagnosed dogs^4^. A minimum score of 16 and a maximum score of 80 were possible, with higher values indicating more severe levels of cognitive dysfunction. Age, sex, sterilization status, breed group, geographic region, physical activity level, comorbidities, and primary formal dog activity were assessed through the HLES.

For owner-reported purebred dogs, breed was elicited and assigned based on the eight groups defined by the American Kennel Club: herding, hound, toy, non-sporting, sporting, terrier, working, and miscellaneous/Foundation Stock Service^18^. Primary formal dog activity refers to a dog’s main job or activity with the owner and was characterized as one of the following: companion animal or pet, obedience, show, breeding, agility, hunting, working, field trials, search and rescue, service dog, or assistance/therapy dog. Physical activity over the past year was classified as not active, moderately active, or very active. Major comorbidities considered were owner-reported in the HLES, including any history of cancer, endocrine disorders, kidney disorders, immune disorders, trauma, toxin consumption, infectious disease, hematologic disorders, neurological disorders, orthopedic disorders, reproductive disorders, liver disorders, gastrointestinal disorders, respiratory disorders, cardiac disorders, skin disorders, oral disorders, eye disorders, or ear disorders. Geographic region was determined based on the owner’s primary state of residence and categorized as West, Southwest, Midwest, Southeast, or Northeast.

### Data Analyses

The primary outcome of interest was total CSLB score. Additionally, canine cognitive function was treated as a binary measure such that dogs with a CSLB score ≥50 were classified as having CCD^4^. Logistic regression models were fit to assess univariable associations between CCD and age, sex, sterilization status, breed group, geographic region, major comorbidities, and activity variables. Covariates that were determined *a priori* to be associated with CCD, informed by a directed acyclic graph (DAG) (Figure 2), were included in a multivariable logistic regression model.

Predicted lifespan quartile was then estimated for each subject. Projected mean life expectancies were calculated on a separate data set collected by Urfer *et al*. consisting of private U.S. veterinary hospital medical records for companion dogs^19^. Mean life expectancies were then assigned to each dog in the DAP data set as a function of their weight or projected weight class, sterilization status, and sex^16,19^. Each dog was assigned a predicted lifespan quartile that was equivalent to the proportion of their current life lived to their projected lifespan. Lifespan quartile was then evaluated for its ability to predict CCD using adjusted and unadjusted logistic regression models. The fitted models were used to construct a Receiver Operating Curve (ROC) for two possible diagnostic thresholds (CSLB score ≥50, representative of previously defined thresholds, and >37, which represented the highest quartile of scores in the data). Finally, area under the curve (AUC) was calculated in order to summarize predictive capacity of the fitted models. All statistical analyses were conducted in RStudio (Version 1.3)^20^. All methods and experimental protocols were carried out in accordance with relevant guidelines and regulations. The University of Washington Institutional Review Board (IRB) reviewed the human subjects involvement for this study and determined that the limited dog owner demographic and lifestyle and anecdotal health information collected in the HLES is human subjects research that meets the qualifications for Exempt Status (Category 2). Given the Exempt status, the IRB did not require informed consent for completion of the surveys, but we nevertheless designed an informed consent form and process that meets the regulatory requirements laid down in U.S. Code of Federal Regulations, Title 45 Department of Health and Human Services Part 46, Protection of Human Subjects.

## Results

Of the 15,019 dogs included in analyses, 19.5% were classified as being in their 4^th^ quartile of life, 24.5% were classified as being in their 3^rd^ quartile of life, 27.0% were in their 2^nd^ quartile of life, and 29.1% of dogs in their 1^st^ quartile of life. A total of 1.4% of dogs were classified as having CCD using the binary cut-off of ≥ 50. There were similar distributions of sex, geographic region, primary role, and breed type among each lifespan quartile (Table 1). Sterilization status, history of each of 17 categories of health problems reported in HLES, purebred breed group, and activity level were substantially positively associated with weight-based projected lifespan quartile. Dogs in a higher weight-based projected lifespan quartile were more likely to be sterilized, more likely to have history of the above comorbidities, and were less active.

**Table 1:**
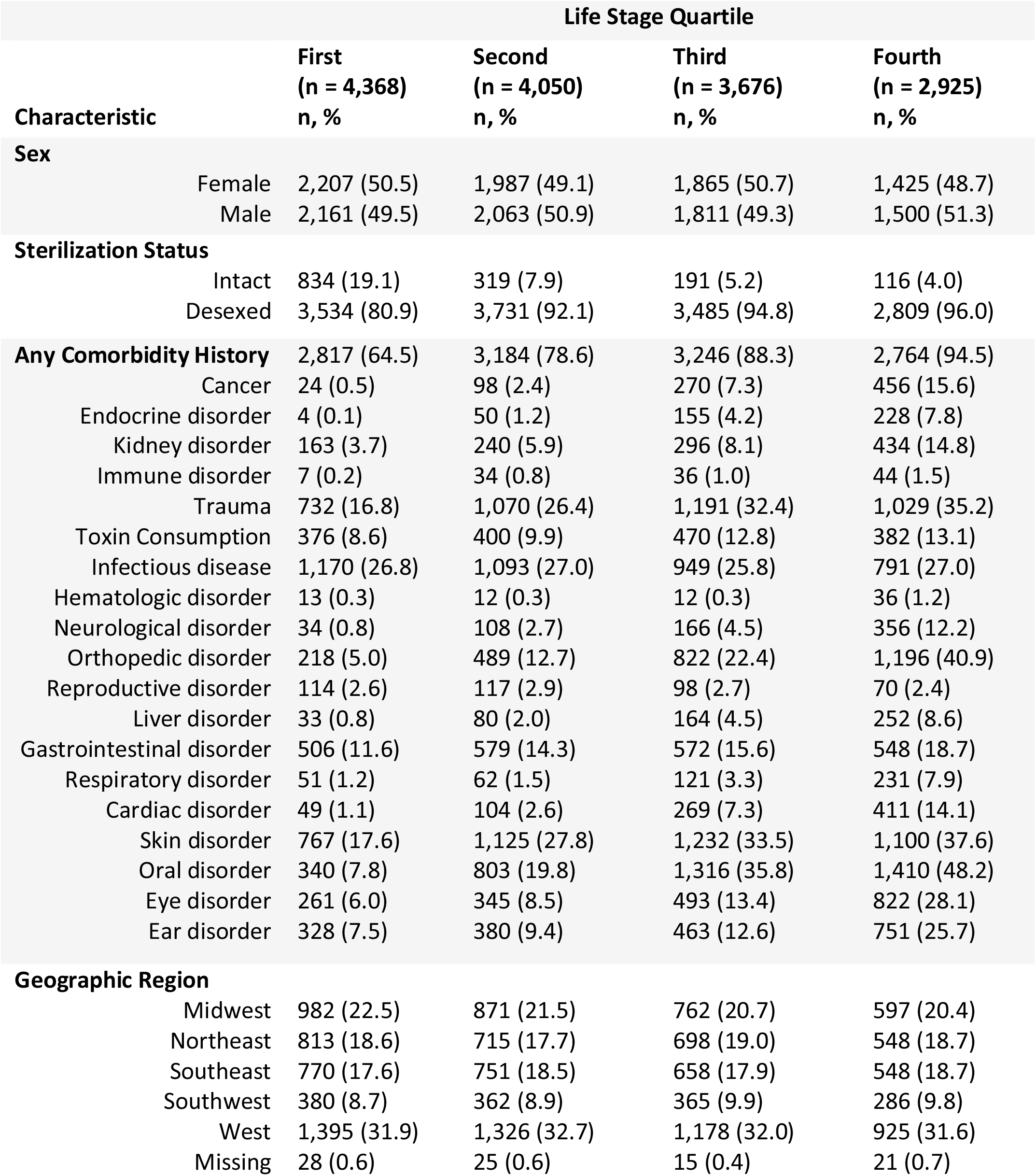
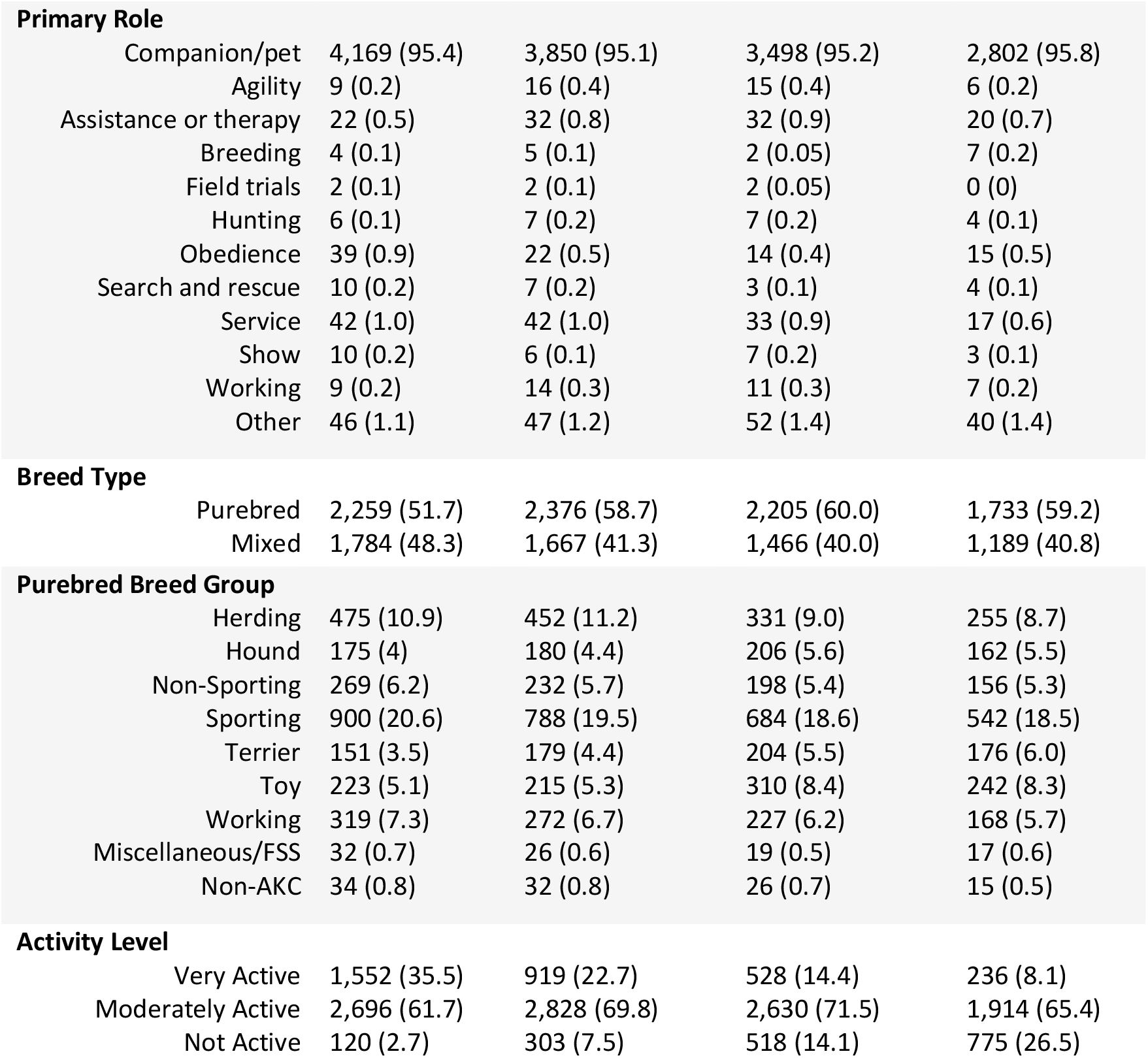
Selected canine demographic characteristics by weight-based projected lifespan quartile (n = 15,019), Dog Aging Project 2020-2021

|  |  | Life Stage Quartile |  |  |  |
| --- | --- | --- | --- | --- | --- |
| Characteristic |  | First<br>(n = 4,368)<br>n, % | Second<br>(n = 4,050)<br>n, % | Third<br>(n = 3,676)<br>n, % | Fourth<br>(n = 2,925)<br>n, % |
| **Sex** |  |  |  |  |  |
|  | Female | 2,207 (50.5) | 1,987 (49.1) | 1,865 (50.7) | 1,425 (48.7) |
|  | Male | 2,161 (49.5) | 2,063 (50.9) | 1,811 (49.3) | 1,500 (51.3) |
| **Sterilization Status** |  |  |  |  |  |
|  | Intact | 834 (19.1) | 319 (7.9) | 191 (5.2) | 116 (4.0) |
|  | Desexed | 3,534 (80.9) | 3,731 (92.1) | 3,485 (94.8) | 2,809 (96.0) |
| **Any Comorbidity History** |  | 2,817 (64.5) | 3,184 (78.6) | 3,246 (88.3) | 2,764 (94.5) |
|  | Cancer | 24 (0.5) | 98 (2.4) | 270 (7.3) | 456 (15.6) |
|  | Endocrine disorder | 4 (0.1) | 50 (1.2) | 155 (4.2) | 228 (7.8) |
|  | Kidney disorder | 163 (3.7) | 240 (5.9) | 296 (8.1) | 434 (14.8) |
|  | Immune disorder | 7 (0.2) | 34 (0.8) | 36 (1.0) | 44 (1.5) |
|  | Trauma | 732 (16.8) | 1,070 (26.4) | 1,191 (32.4) | 1,029 (35.2) |
|  | Toxin Consumption | 376 (8.6) | 400 (9.9) | 470 (12.8) | 382 (13.1) |
|  | Infectious disease | 1,170 (26.8) | 1,093 (27.0) | 949 (25.8) | 791 (27.0) |
|  | Hematologic disorder | 13 (0.3) | 12 (0.3) | 12 (0.3) | 36 (1.2) |
|  | Neurological disorder | 34 (0.8) | 108 (2.7) | 166 (4.5) | 356 (12.2) |
|  | Orthopedic disorder | 218 (5.0) | 489 (12.7) | 822 (22.4) | 1,196 (40.9) |
|  | Reproductive disorder | 114 (2.6) | 117 (2.9) | 98 (2.7) | 70 (2.4) |
|  | Liver disorder | 33 (0.8) | 80 (2.0) | 164 (4.5) | 252 (8.6) |
|  | Gastrointestinal disorder | 506 (11.6) | 579 (14.3) | 572 (15.6) | 548 (18.7) |
|  | Respiratory disorder | 51 (1.2) | 62 (1.5) | 121 (3.3) | 231 (7.9) |
|  | Cardiac disorder | 49 (1.1) | 104 (2.6) | 269 (7.3) | 411 (14.1) |
|  | Skin disorder | 767 (17.6) | 1,125 (27.8) | 1,232 (33.5) | 1,100 (37.6) |
|  | Oral disorder | 340 (7.8) | 803 (19.8) | 1,316 (35.8) | 1,410 (48.2) |
|  | Eye disorder | 261 (6.0) | 345 (8.5) | 493 (13.4) | 822 (28.1) |
|  | Ear disorder | 328 (7.5) | 380 (9.4) | 463 (12.6) | 751 (25.7) |
| **Geographic Region** |  |  |  |  |  |
|  | Midwest | 982 (22.5) | 871 (21.5) | 762 (20.7) | 597 (20.4) |
|  | Northeast | 813 (18.6) | 715 (17.7) | 698 (19.0) | 548 (18.7) |
|  | Southeast | 770 (17.6) | 751 (18.5) | 658 (17.9) | 548 (18.7) |
|  | Southwest | 380 (8.7) | 362 (8.9) | 365 (9.9) | 286 (9.8) |
|  | West | 1,395 (31.9) | 1,326 (32.7) | 1,178 (32.0) | 925 (31.6) |
|  | Missing | 28 (0.6) | 25 (0.6) | 15 (0.4) | 21 (0.7) |

|  |  |  |  |  |
| --- | --- | --- | --- | --- |
| **Primary Role** |  |  |  |  |
| Companion/pet | 4,169 (95.4) | 3,850 (95.1) | 3,498 (95.2) | 2,802 (95.8) |
| Agility | 9 (0.2) | 16 (0.4) | 15 (0.4) | 6 (0.2) |
| Assistance or therapy | 22 (0.5) | 32 (0.8) | 32 (0.9) | 20 (0.7) |
| Breeding | 4 (0.1) | 5 (0.1) | 2 (0.05) | 7 (0.2) |
| Field trials | 2 (0.1) | 2 (0.1) | 2 (0.05) | 0 (0) |
| Hunting | 6 (0.1) | 7 (0.2) | 7 (0.2) | 4 (0.1) |
| Obedience | 39 (0.9) | 22 (0.5) | 14 (0.4) | 15 (0.5) |
| Search and rescue | 10 (0.2) | 7 (0.2) | 3 (0.1) | 4 (0.1) |
| Service | 42 (1.0) | 42 (1.0) | 33 (0.9) | 17 (0.6) |
| Show | 10 (0.2) | 6 (0.1) | 7 (0.2) | 3 (0.1) |
| Working | 9 (0.2) | 14 (0.3) | 11 (0.3) | 7 (0.2) |
| Other | 46 (1.1) | 47 (1.2) | 52 (1.4) | 40 (1.4) |
| **Breed Type** |  |  |  |  |
| Purebred | 2,259 (51.7) | 2,376 (58.7) | 2,205 (60.0) | 1,733 (59.2) |
| Mixed | 1,784 (48.3) | 1,667 (41.3) | 1,466 (40.0) | 1,189 (40.8) |
| **Purebred Breed Group** |  |  |  |  |
| Herding | 475 (10.9) | 452 (11.2) | 331 (9.0) | 255 (8.7) |
| Hound | 175 (4) | 180 (4.4) | 206 (5.6) | 162 (5.5) |
| Non-Sporting | 269 (6.2) | 232 (5.7) | 198 (5.4) | 156 (5.3) |
| Sporting | 900 (20.6) | 788 (19.5) | 684 (18.6) | 542 (18.5) |
| Terrier | 151 (3.5) | 179 (4.4) | 204 (5.5) | 176 (6.0) |
| Toy | 223 (5.1) | 215 (5.3) | 310 (8.4) | 242 (8.3) |
| Working | 319 (7.3) | 272 (6.7) | 227 (6.2) | 168 (5.7) |
| Miscellaneous/FSS | 32 (0.7) | 26 (0.6) | 19 (0.5) | 17 (0.6) |
| Non-AKC | 34 (0.8) | 32 (0.8) | 26 (0.7) | 15 (0.5) |
| **Activity Level** |  |  |  |  |
| Very Active | 1,552 (35.5) | 919 (22.7) | 528 (14.4) | 236 (8.1) |
| Moderately Active | 2,696 (61.7) | 2,828 (69.8) | 2,630 (71.5) | 1,914 (65.4) |
| Not Active | 120 (2.7) | 303 (7.5) | 518 (14.1) | 775 (26.5) |

Associations between various canine characteristics and binary CCD assignment were assessed using univariable and multivariable logistic regression models (Table 2). When assessing age alone, the odds of being diagnosed with CCD increased almost 70% with each additional year of age (OR=1.68, 95% CI 1.60, 1.77) (Figure 1).

**Table 2:** Association between selected dog characteristics and Canine Cognitive Dysfunction, Dog Aging Project 2020-2021

| Characteristic (n) | Univariable Analysis |  | Multivariable Analysis |  |
| --- | --- | --- | --- | --- |
|  | OR | 95% CI | OR | 95% CI |
| **Age (integer years)** | 1.68 | 1.60-1.77 | 1.52 | 1.44-1.61 |
| **Sex** |  |  |  |  |
| Female (reference) (7,484) | 1.00 | - | - | - |
| Male (7,535) | 0.91 | 0.69-1.19 |  |  |
| **Sterilization status** |  |  |  |  |
| Desexed (reference) (13,559) | 1.00 | - | 1.00 | - |
| Intact (1,460) | 0.36 | 0.16-0.67 | 1.21 | 0.51-2.51 |
| **Activity level** |  |  |  |  |
| Very Active (reference) (3,235) | 1.00 | - | 1.00 | - |
| Moderately Active (10,068) | 4.80 | 2.29-12.33 | 2.19 | 1.00-5.83 |
| Not Active (1,716) | 40.46 | 19.42-103.51 | 6.47 | 2.93-17.23 |
| **Breed group** |  |  |  |  |
| Sporting (reference) (2,914) | 1.00 | - |  |  |
| Herding (1,513) | 1.07 | 0.55-1.99 |  |  |
| Hound (723) | 1.65 | 0.78-3.26 |  |  |
| Terrier (710) | 3.58 | 2.02-6.28 |  |  |
| Toy (990) | 3.80 | 2.29-6.38 | - | - |
| Non-Sporting (855) | 3.49 | 2.03-6.00 |  |  |
| Working (986) | 0.76 | 0.31-1.67 |  |  |
| Misc./FSS (94) | 0.00 | - |  |  |
| Non-AKC (107) | 1.01 | 0.05-4.80 |  |  |
| **Geographic region** |  |  |  |  |
| West (reference) (4,824) | 1.00 | - |  |  |
| Midwest (3,212) | 0.82 | 0.55-1.19 |  |  |
| Northeast (2,774) | 0.90 | 0.61-1.32 | - | - |
| Southeast (2,727) | 0.82 | 0.55-1.22 |  |  |
| Southwest (1,393) | 0.81 | 0.47-1.32 |  |  |
| **History of Cancer** |  |  |  |  |
| No (reference) (14,171) | 1.00 | - | 1.00 | - |
| Yes (848) | 4.09 | 2.85-5.72 | 1.35 | 0.90-1.97 |
| **History of Kidney disorder** |  |  |  |  |
| No (reference) (13,886) | 1.00 | - | 1.00 | - |
| Yes (1,133) | 3.53 | 2.52-4.87 | 1.21 | 0.82-1.76 |
| **History of Endocrine disorder** |  |  |  |  |
| No (reference) (14,582) | 1.00 | - | 1.00 | - |
| Yes (437) | 3.74 | 2.30-5.80 | 1.09 | 0.63-1.79 |
| **History of Immune disorder** |  |  |  |  |
| No (reference) (14,898) | 1.00 | - | - | - |
| Yes (121) | 1.16 | 0.19-3.67 |  |  |
| **History of Trauma** |  |  |  |  |
| No (reference) (10,997) | 1.00 | - | 1.00 | - |
| Yes (4,022) | 1.63 | 1.23-2.15 | 1.10 | 0.79-1.50 |
| **History of Toxin Consumption** |  |  |  |  |
| No (reference) (13,391) | 1.00 | - | 1.00 | - |
| Yes (1,628) | 1.61 | 1.10-2.29 | 1.13 | 0.73-1.69 |
| **History of Infectious disease** |  |  |  |  |
| No (reference) (11,016) | 1.00 | - | - | - |
| Yes (4,003) | 0.81 | 0.58-1.11 |  |  |
| **History of hematologic disorder** |  |  |  |  |
| No (reference) (14,946) | 1.00 | - | - | - |
| Yes (73) | 5.16 | 1.79-11.71 |  |  |
| **History of Neurological disorder** |  |  |  |  |
| No (reference) (14,355) | 1.00 | - | 1.00 | - |
| Yes (664) | 8.22 | 5.96-11.20 | 1.84 | 1.26-2.65 |
| **History of Orthopedic disorder** |  |  |  |  |
| No (reference) (12,294) | 1.00 | - | 1.00 | - |
| Yes (2,725) | 3.33 | 2.52-4.37 | 0.88 | 0.64-1.20 |
| **History of Reproductive disorder** |  |  |  |  |
| No (reference) (14,620) | 1.00 | - | - | - |
| Yes (399) | 0.69 | 0.21-1.64 |  |  |
| **History of Liver disorder** |  |  |  |  |
| No (reference) (14,490) | 1.00 | - | 1.00 | - |
| Yes (529) | 3.38 | 2.12-5.15 | 0.96 | 0.57-1.57 |
| **History of Gastrointestinal disorder** |  |  |  |  |
| No (reference) (12,814) |  |  |  |  |
| Yes (2,205) | 1.00 | - | 1.00 | - |
|  | 1.59 | 1.13-2.19 | 1.10 | 0.76-1.58 |
| **History of Respiratory disorder** |  |  |  |  |
| No (reference) (14,554) | 1.00 | - | 1.00 | - |
| Yes (465) | 3.69 | 2.29-5.67 | 0.75 | 0.44-1.22 |
| **History of Cardiac disorder** |  |  |  |  |
| **No (reference) (14,186)** | 1.00 | - | 1.00 | - |
| **Yes (833)** | 3.78 | 2.61-5.35 | 0.83 | 0.54-1.24 |
| **History of Skin disorder** |  |  |  |  |
| **No (reference) (10,795)** | 1.00 | - | 1.00 | - |
| **Yes (4,224)** | 1.68 | 1.27-2.21 | 0.84 | 0.61-1.15 |
| **History of Oral disorder** |  |  |  |  |
| **No (reference) (11,150)** | 1.00 | - | 1.00 | - |
| **Yes (3,869)** | 3.58 | 2.74-4.71 | 0.79 | 0.58-1.08 |
| **History of Eye disorder** |  |  |  |  |
| **No (reference) (13,098)** | 1.00 | - | 1.00 | - |
| **Yes (1,921)** | 8.94 | 6.81-11.78 | 2.16 | 1.57-2.98 |
| **History of Ear disorder** |  |  |  |  |
| **No (reference) (13,046)** | 1.00 | - | 1.00 | - |
| **Yes (1,973)** | 6.56 | 5.00-8.62 | 1.96 | 1.42-2.70 |
| **Breed** |  |  |  |  |
| **Mixed (reference) (6,106)** | 1.00 | - | 1.00 | - |
| **Purebred (8,913)** | 1.43 | 1.07-1.91 | 1.24 | 0.90-1.72 |
| **Primary activity** |  |  |  |  |
| **Pet (reference) (14,319)** | 1.00 | - |  |  |
| **Agility (46)** | 1.49 | 0.08-6.85 |  |  |
| **Assistance or therapy (106)** | 0.00 | - |  |  |
| **Breeding (18)** | 0.00 | - |  |  |
| **Field Trials (6)** | 0.00 | - |  |  |
| **Hunting (24)** | 0.00 | - | - | - |
| **Obedience (90)** | 0.00 | - |  |  |
| **Other (185)** | 0.73 | 0.12-2.30 |  |  |
| **Search and Rescue (24)** | 0.00 | - |  |  |
| **Service (134)** | 0.50 | 0.03-2.26 |  |  |
| **Show (26)** | 0.00 | - |  |  |
| **Working (41)** | 0.00 | - |  |  |

**Figure 1:**
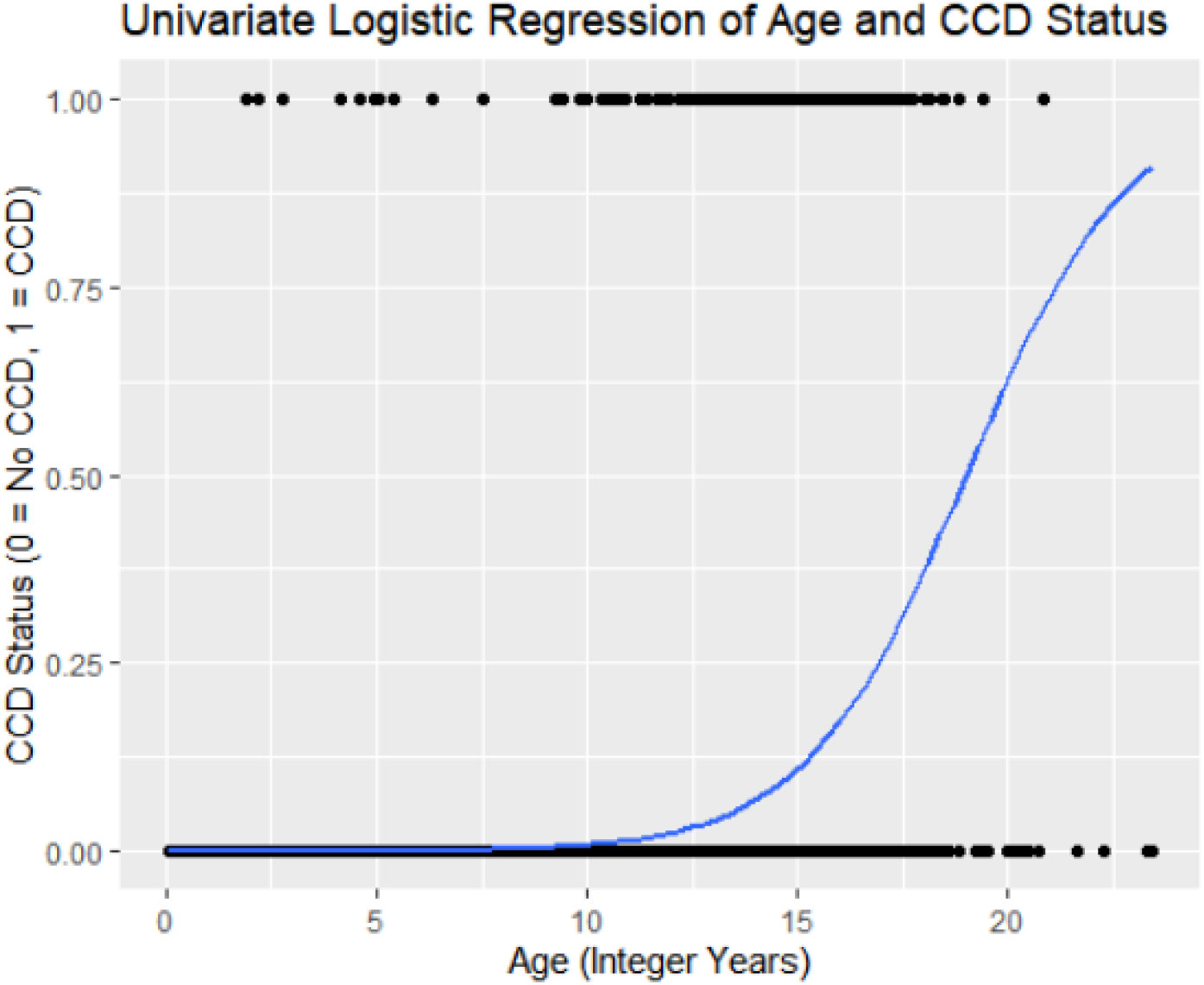
Logistic Regression curve representing association between age and CCD status, Dog Aging Project, 2020-2021

A directed acyclic graph (DAG) was constructed to highlight relevant covariates to include in the multivariable logistic regression model assessing the association between age and CCD (Figure 2). Age, sterilization status, history of 15 categories of health problems, breed type, and activity level were selected for the final logistic regression model (Table 2). When controlling for all other characteristics, the odds of CCD increased 52% with each additional year of age (OR=1.52, 95% CI 1.44, 1.61). Interestingly, among dogs of a given age, health status, breed type, and activity level, those who were intact had a higher odds of CCD (OR=1.21, 95% CI 0.51-2.51). Although attenuated, an inverse association between activity level and CCD was still found. Among dogs of the same age, health status, breed type, and sterilization status, odds of CCD were 6.47 times higher in dogs who were not active compared to those who were very active (OR=6.47, 95% CI 2.93-17.23). When controlling for age, breed type, activity level, and other comorbidities, dogs with a history of neurological, eye, or ear disorders had higher odds of CCD (OR=1.84, 95% CI 1.26, 2.65; OR=2.16, 95% CI 1.57, 2.98; OR=1.96, 95% 1.42, 2.70, respectively).

**Figure 2:**
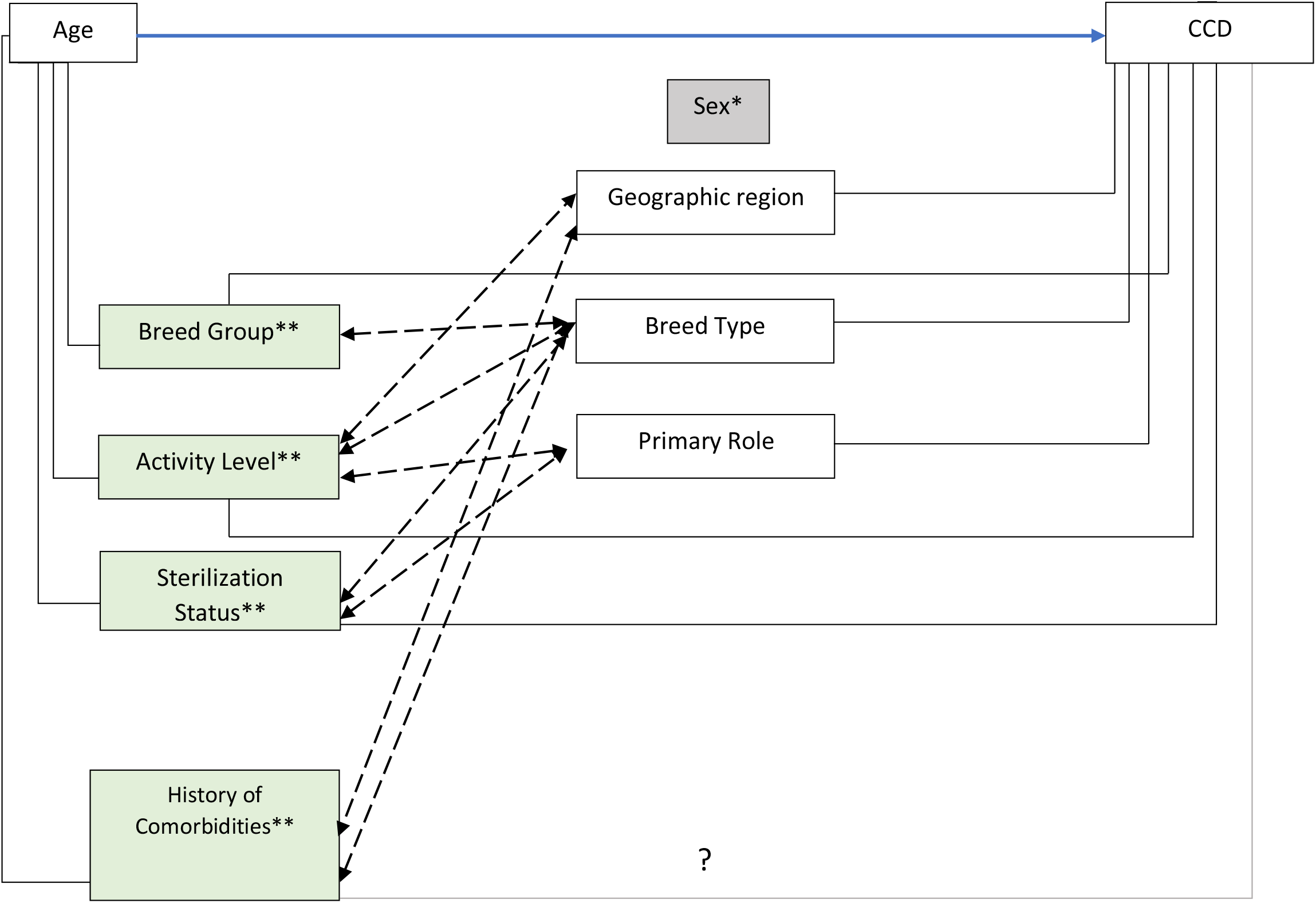
Directed Acyclic Graph (DAG) representing associations between Canine Cognitive Dysfunction, age, and other characteristics of interest, Dog Aging Project, 2020-2021 *Sex was not associated with either age or CCD, and so was not included in the DAG. **These variables were identified as confounders that were associated with age and CCD and were included in the multivariable models. Dashed lines represent hypothesized associations considered based on relevance

Adjusted and unadjusted logistic regression models were then fit using the weight-based projected lifespan quartiles. Adjusted models included all covariates from the multivariable logistic regression model assessing the association between age and CCD (Table 2). These weight-based projected lifespan quartile models were evaluated for their ability to accurately classify CCD at two different diagnostic thresholds: ≥ 50, as cited in previous literature, and > 37, which captures the upper 4^th^ quartile of CSLB scores. In total, four predictive models were assessed by evaluating the area under the curve (AUC) of an ROC curve (Figure 3). The diagnostic threshold of CSLB score ≥ 50 had the highest predictive capacity. The adjusted model that controlled for health status, breed type, sterilization status, and activity level provided a slightly improved predictive capacity over the unadjusted model (AUC=0.892, 95% CI: 0.854-0.930 vs. AUC=0.862, 95% CI: 0.831-0.893). The threshold of CSLB score > 37 had lower predictive capacity for both the adjusted and unadjusted models (AUC=0.701, 95% CI: 0.682-0.721 vs. AUC=0.644, 95% CI: 0.624-0.665).

**Figure 3:**
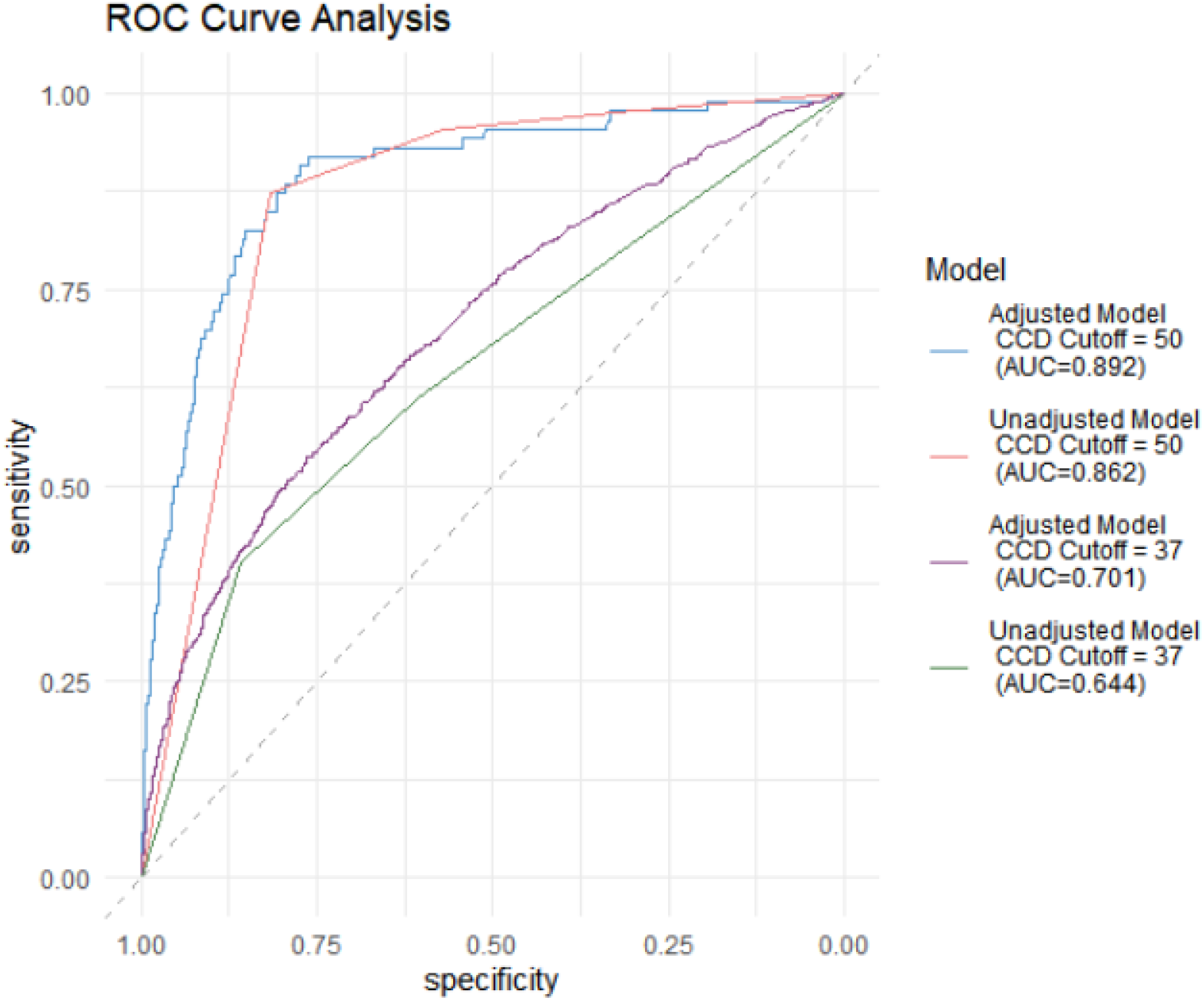
ROC curve generated from predictive models relating selected dog characteristics to Canine Cognitive Dysfunction prevalence, Dog Aging Project 2020-2021

## Discussion

Results from this study indicate a positive association between age and CCD in companion dogs, even adjusting for other characteristics in a multivariable logistic regression model. There also existed associations between CCD and decreased activity level, as well as CCD and history of an eye, ear, or neurological disorder. The association between age and CCD aligns with the progressive nature of CCD and with previous dog research findings, which have shown an exponential increase in CCD prevalence with increasing age^4^.

An inverse association was seen in dogs whose owners indicated higher dog activity levels over the past year. Previous studies with rodent models have demonstrated that exercise can have protective effects against the development of biological markers and subsequent behavioral deficits characteristic of AD, and numerous observational human studies have consistently shown inverse associations between exercise and AD^21, 22, 23, 24, 25^. These observations may reflect a variety of biologic mechanisms, including a reduction of pro-inflammatory cytokines in the brain that otherwise contribute to neural damage and death, and an increase in neural plasticity^24,25^. The reduced odds of CCD among more active dogs in our cohort may be a result of these same mechanisms, but it is important to note that this correlation could also exist simply because of dogs exhibiting less activity due to their cognitive decline.

The association we observed between dogs who have ever had a neurological disorder and their prevalence of CCD is expected. Dementia, a neurological disorder that could have been indicated by an owner in their HLES survey, is characterized by a decline in cognitive function. Seizure disorders, which also could have been indicated by the owner, have been found to occur more commonly in human AD patients than in comparable individuals without AD^26^. Human studies have demonstrated potential links between eye and ear disorders and AD. Impairments in visual acuity and visual fields have been observed more frequently in AD patients than in similarly aged individuals without AD^27,28^. Further, the presence of ophthalmic conditions, such as cataracts and age-related macular degeneration, has been found to be associated with an increased risk of all-cause dementia and AD^29^. Amyloid deposits, which have been linked to the development of AD and have been associated with the acceleration of AD progression, have been found in the lens of the eye in some individuals with age-related cataracts, potentially indicating common neuropathological pathways leading to AD and cataracts^30^. Age-related macular degeneration has also been associated with an increased risk of all-cause dementia and AD^30^. Like the potential relationship between cataracts and AD, amyloid deposits have been detected in the small fatty protein deposits that accumulate under the eye of individuals with age-related macular degeneration^31^.

In addition to numerous studies identifying hearing loss as a possible risk factor for cognitive decline and all-cause dementia, a recent study found that individuals with dual sensory impairment (combined visual and hearing impairments) had a substantially increased risk of AD, possibly as a result of reduced neural resources needed for cognitive performance^31, 32, 33, 34.^ These associations are potentially mirrored in our companion dog cohort.

Alternatively, the associations we observed between sensory impairment and CCD could be due at least in part to misclassification. The CSLB survey, which is closely derived from Salvin *et al*.’s CCDR, is intended to distinguish between behaviors associated with CCD as opposed to those that are part of normal dog aging^4^. However, some behaviors assessed by the rating scale may not arise solely from cognitive impairment^4^. Questions such as, “ How often does your dog walk into walls or doors?” and “ How often does your dog have difficulty finding food dropped on the floor?” were included in the CCDR because increased frequency of these behaviors is seen in dogs with CCD. However, sensory impairments could also cause increased frequency of these behaviors, resulting in high scores on the instrument.

We observed a dog’s estimated lifespan quartile, which is a function of their age, weight, sex, and sterilization status, to have excellent discriminating ability between CCD positive and negative dogs (CSLB score ≥ 50). This improved only slightly in the model adjusted for age, sterilization status, history of 15 categories of health problems, breed type, and activity level. We also considered a lower diagnostic threshold for CCD assignment by using the top quartile of CSLB scores as a cut-off. However, that classification yielded much lower discriminating ability between CCD positive and negative dogs for lifespan quartile. Although the diagnostic threshold of 50 had high predictive capacity, it is important to note that this threshold between normal aging and CCD was chosen by previous researchers, and measures of sensitivity and specificity would be difficult to obtain^4^. It would therefore be beneficial to examine the validity of this diagnostic tool more closely in future studies.

This study had several strengths, including a large sample size, standardized data collection methods, high participant response proportions within the cohort, and the inclusion of many potential confounders in analytic models. Important limitations did exist in this study that are worth addressing. In addition to potential concerns with the sensitivity and specificity of the diagnostic threshold chosen for CCD, that this analysis considers only the first year of data and is not yet longitudinal, it is not possible to assess any prospective links between the various canine characteristics and CCD. While the HLES and CSLB surveys were administered to owners at different times (the maximum time interval between the completion of the HLES and CSLB surveys was 50.3 weeks), the median time between survey completion was only 5 weeks. 11,324 of the total 15,019 owners completed both surveys within 10 weeks. There was a low likelihood that this timeframe would result in any sort of meaningful changes in CCD status. Future studies using prospective DAP data will be able to explore potential risk factors for CCD as well as cognitive decline over time.

Another limitation is that all data used in this analysis were based on surveys completed by owners. Owner-reported information is potentially susceptible to multiple forms of bias. Questions requiring owners to recall behaviors and medical conditions that could have occurred from 6 months to many years prior could introduce either differential or non-differential misclassification, depending on whether the errors in recall were correlated with the dog’s cognitive status. Non-differential misclassification could arise from social desirability if, for example, owners indicated a higher level of physical activity for their dog, or a lower level of impairment on the CSLB survey. Such errors would tend to dampen associations between physical activity and CCD.

It is also important to recognize the drastic changes that have taken place in many households due to the COVID-19 pandemic, and the possibility that these changes influenced our results. Depending on when an owner completed their HLES and CSLB surveys, their dog’s activity level may have changed as a result of stay-at-home orders and/or the owner’s ability to work from home. Additionally, owners spending more time at home with their dog may have an impact on the dog’s health, and it may increase the likelihood of observing specific health behaviors that would affect a dog’s reported health status. These types of major lifestyle changes could have altered our data in ways that would be difficult to quantify. Finally, there is the potential for unmeasured confounders that were not captured by our surveys, as well as survey data that were not considered for analysis, which could result in residual confounding potentially adding bias to our results.

We identified a positive association between companion dog age and CCD. We also identified strong positive associations between history of a neurological, ear, or eye disorder as well as an association between decreased physical activity level and having a CSLB score of 50 or higher. Dogs were classified into their predicted quartile of life based on their age, weight, sex, and sterilization status. Using a binary diagnostic threshold of ≥50, a dog’s lifespan quartile showed excellent discriminating ability between CCD positive and negative dogs. This quartile estimation could potentially serve as a useful tool to inform whether a dog should be screened for CCD by their veterinarian. Finally, given increasing evidence of the parallels between canine and human cognitive disease, accurate CCD diagnosis in dogs may provide researchers with more suitable animal models in which to study aging in human populations.

## Data Availability Statement

The datasets generated during and/or analyzed during the current study are available through Terra in the Dog Aging Project 2020 Curated Workspace.

## Author Contributions

SY performed statistical analysis and contributed to the drafting and editing of the manuscript. AF, SMS, KC, and AR provided statistical advice and contributed to the drafting and editing of the manuscript. VW, MK, and DP provided scientific advice and contributed to the drafting and editing of the manuscript. The authors read and approved the final manuscript.

## Additional Information

### Dog Aging Project Consortium

Kate Creevy^3^, Audrey Ruple^4^, Vanessa Wilkins^3^, Matt Kaeberlein^5^, Daniel Promislow^6^

^3^ College of Veterinary Medicine & Biomedical Sciences, Texas A&M University, College Station TX

^4^ Department of Population Health Sciences, Virginia-Maryland College of Veterinary Medicine, Virginia Tech, Blacksburg VA

^5^ Department of Laboratory Medicine & Pathology, University of Washington School of Medicine, Seattle WA

^6^ Department of Biology, University of Washington, Seattle WA

